# PINK1 deficiency rewires early immune responses in a mouse model of Parkinson’s disease triggered by intestinal infection

**DOI:** 10.1101/2024.06.18.598931

**Authors:** Sherilyn Junelle Recinto, Alexandra Kazanova, Hicham Bessaiah, Brendan Cordeiro, Sriparna Mukherjee, Shobina Premachandran, Adam MacDonald, Jessica Pei, Alexis Allot, Elia Afanasiev, Moein Yaqubi, Heidi M McBride, Louis-Eric Trudeau, Samantha Gruenheid, Jo Anne Stratton

## Abstract

Parkinson’s disease (PD) is characterized by a protracted period of non-motor symptoms, including gastrointestinal (GI) dysfunction, which can precede the development of the cardinal motor deficits by decades. This long prodrome of disease is highly suggestive of immune cell involvement in the initiation of disease, but currently the field lacks robust model systems to study such mechanisms. It has been hypothesized that pathology may be first initiated in the periphery due to environmental triggers, such as pathogens that enter the GI tract. We further speculate that the impact of such pathogens on the immune system could be exacerbated in genetically predisposed individuals. Our group has developed a GI-targeted pathogen-induced PD mouse model system in PINK1 KO mice with Gram-negative bacterial infections and found that T cells are a major player in driving PD-like motor symptoms at late stages following infection. Herein, we now map the initiating immune events at the site of infection at the earliest stages with the goal of shedding light on the earliest mechanisms triggering immune-mediated pathological processes relevant to PD. Using unbiased single cell sequencing, we demonstrate that myeloid cells are the earliest dysregulated immune cell type in PINK1 KO infected mice at 1-week post-infection, followed by a dysregulated T cell response shortly after, at 2 weeks post-infection. We find that these myeloid cells have an enhanced proinflammatory profile, are more mature, and develop enhanced capacity for antigen presentation. Using unbiased prediction analysis, our data suggest that cytotoxic T cells and myeloid cells are particularly poised for interacting with each other, and we identify possible direct cell-cell interaction pathways that might be implicated. Taken together, deciphering the earliest immune mechanisms in the periphery underpinning PD autoimmunity will be instrumental in the development of effective therapeutic targeting strategies before irrevocable neuronal damage ensues.

## Introduction

Parkinson’s disease (PD) is a highly prevalent, age-associated progressive neurodegenerative disorder that is usually diagnosed upon the appearance of cardinal motor functional deficits such as bradykinesia, tremor, and rigidity. The motor symptoms are caused in part by the loss of dopamine (DA) neurons in the substantia nigra pars compacta of the midbrain and it is estimated that by the time of diagnosis, 60-80% of nigral DA neurons are already irrevocably lost, making treatment of PD challenging [1]. Nevertheless, it is now widely recognized that motor symptom development can be preceded by a long prodromal phase where non-motor symptoms, such as hyposmia, constipation, sleep disruption and anxiety, may be present [2]. The extensive prodromal period suggests that disease initiating mechanisms may occur early life and remain unnoticed until the appearance of motor symptoms in older age. Understanding the pathophysiology of the prodromal phase of PD offers promise for earlier detection and treatment.

Peripheral inflammation and immune dysfunction have been implicated in the PD onset, including the emergence of various non-motor symptoms, such as gastrointestinal (GI) dysfunction [3]. Analyses of immune profiles in the stools of PD patients demonstrate higher levels of inflammatory mediators suggestive of GI inflammation [4-8]. Clinical studies further reveal that people with inflammatory bowel disease (IBD) and GI infections have an increased risk of developing PD later in life [4, 9-13]. Other studies also underscored common genetic variants between patients with PD and autoimmune disorders, such as IBD [14, 15]. Intriguingly, converging evidence suggests that multiple genes implicated in familial forms of PD as well as several identified PD risk variants are expressed in immune cells and function in the regulation of immune responses.

Of interest, *PTEN-induced kinase 1* (PINK1), a serine/threonine kinase, and Parkin (PRKN), an E3 ubiquitin ligase, both associated with familial forms of PD, have been shown to act as repressors of innate immune responses independently from their previously described roles in mitophagy. A loss of function of either of these two proteins disinhibits the formation of mitochondrial-derived vesicles (MDVs) in antigen-presenting cells (APCs) leading to the presentation of mitochondrial peptides via major histocompatibility complex I (MHCI), a process referred to as mitochondrial antigen presentation (MitAP) [16]. MitAP leads to the establishment of peripheral mitochondrial antigen-specific T cells, implicating T cell autoimmunity in PD pathogenesis. Subsequently, others have also proposed that PINK1 modulates innate immune responses triggered by oxidative stress and viral infection [17-20].

The occurrence of MitAP in the periphery raises the notion that peripheral mitochondrial antigen-specific T cells may access the brain, for example, in the context of infections. In accordance with this hypothesis, intestinal infection with Gram-negative bacteria (*Citrobacter rodentium*) in PINK1 KO mice led to antigen-specific T-cell autoreactivity and increased entry of immune cells into the brain [21]. Notably, with aging and repeated infections, these mice developed L-DOPA reversible PD-like motor impairments. A similar phenomenon was also recently demonstrated using a major PD-associated human-relevant pathogen (*Helicobacter pylori*), where autoreactive CD8+ T-dependent PD-like motor phenotypes developed following GI infection of PINK1 KO, but not wild type littermate mice [22]. Additionally, adoptive transfer of mitochondrial antigen-specific CD8+ T cells was recently shown to be sufficient to induce motor impairments and DA neuron loss in both PINK1 KO and wild-type mice, supporting a critical and direct role for autoreactive CD8+ T cells in causing neuronal loss under such conditions [23]. Together, these data provide proof of concept that peripheral infections can lead to aberrant immunity engendering parkinsonism in genetically predisposed models of PD.

Critical questions remain, in particular regarding the mechanisms at play in the periphery at the earliest prodromal stages of the disease process. Understanding the immune mechanisms that initiate disease early in the periphery at the cell/molecular level will address a major knowledge gap that hinders the development of diagnostic and therapeutic approaches to stop the onset of motor impairment at early stages of the disease before irreversible neuron death occurs. Here, we aimed to provide an in-depth characterisation of the initiating immunological events in the gut mediated by PINK1 deficiency. We find that intestinal infection in PINK1 KO mice induces early immune dysregulation in the colon marked by the promotion of proinflammatory myeloid cell differentiation and an increased capacity for antigen presentation. We also find an expansion of intestinal CD8+ T cells, which we postulate is due to mis-presentation of self-peptides, laying the foundation for autoimmunity later in life. Gaining insight into the role of PD-associated genes and mutations in the dysregulation of the early immune responses in the periphery provides an opportunity for the development of better biomarkers and effective therapeutic targeting prior to the occurrence of irrevocable neuronal damage that denotes the later stages of PD.

## Results

### Single cell RNAseq of the colon unveils a role of PINK1 within the immune cell compartment at the earliest stages following infection

We have previously described that repeated infection of PINK1 KO mice with the mouse intestinal Gram-negative pathogen *C. rodentium* via gavage induces L-DOPA reversible motor symptoms 16-to 24-weeks later [21]. Despite clear evidence for the role of autoreactive CD8+ T cells in the neurodegenerative process within the brain [22, 23], it is unknown how this is instigated. The phases of *C. rodentium* infection subsequent to its establishment in the colon are well described, and include bacterial expansion (by 1 week), steady-state (1-2 weeks) and clearance (2+ weeks). This immune response is governed by several players of innate and adaptive immunity, including early myeloid cell responses driven by dendritic cells then ultimately the engagement of type I and interleukin 17-producing T helper cells (Th1 and Th17, respectively) for complete clearance of the infection [24].

We first employed an indirect, non-invasive approach using fecal analysis to evaluate whether there was any evidence for enhanced peripheral inflammation in PINK1 KO mice at different time points after infection. We used enzyme-linked immunosorbent assay (ELISA) to detect lipocalin-2 (LCN2), which is highly expressed by activated innate immune/epithelial cells, and often used as an indicator of intestinal inflammation [25, 26] (Fig. 1A). We noted a progressive increase in LCN2 concentration over time which reached significance in wild type and PINK1 KO mice at 2- and 4-w.p.i. compared to uninfected mice. Importantly, there was also significantly more LCN2 in PINK1 KO infected mice compared to wild type infected mice at these time points.

**Figure 1:**
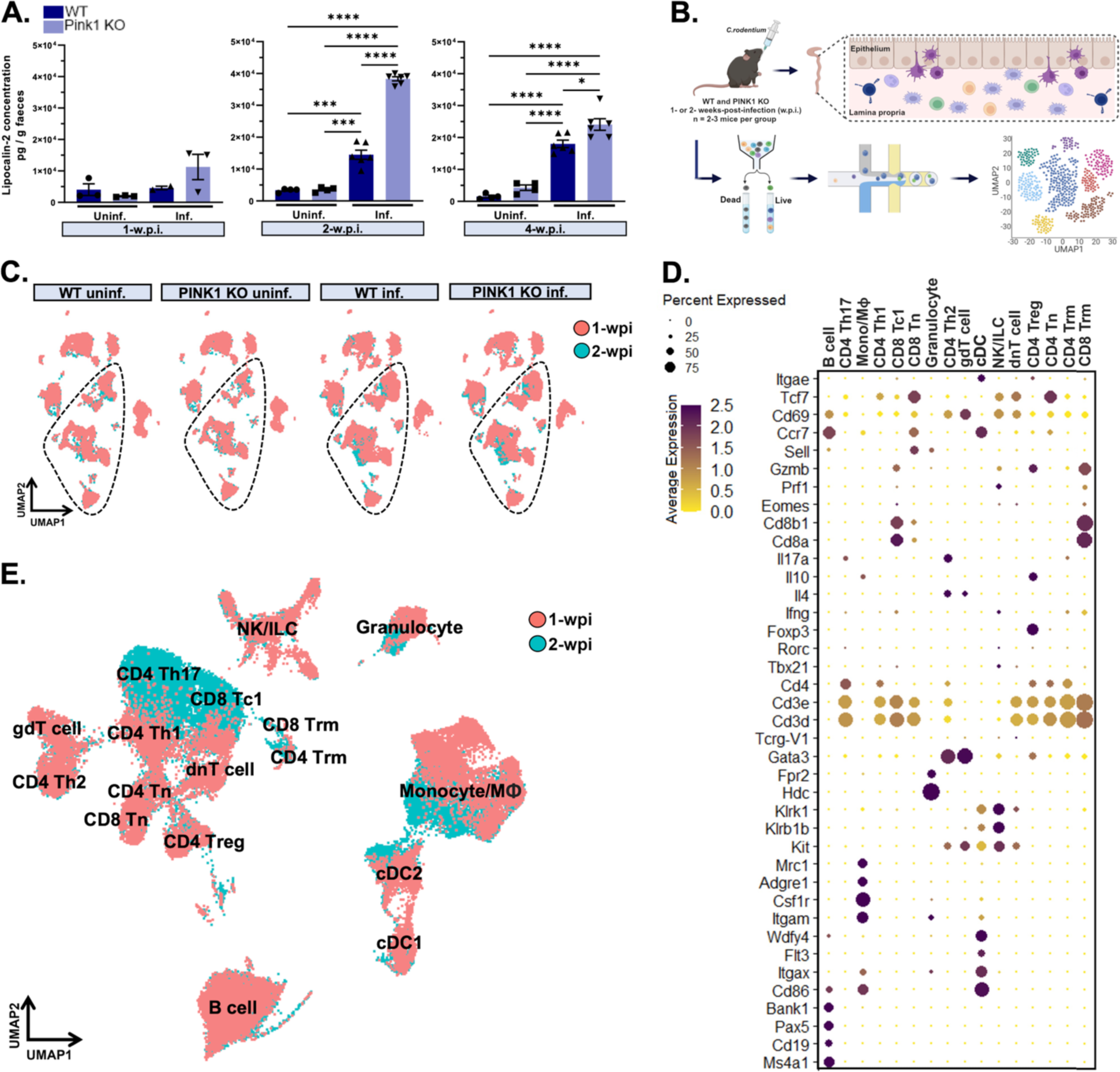
Single-cell transcriptomics of the gut in an infection-induced model of Parkinson’s disease. A) Lipocalin-2 levels were measured in fecal samples of WT and PINK1 KO mice at 1-, 2- and 4- w.p.i. and uninf. controls. Two-way ANOVA followed by Tukey’s multiple comparison test was applied. Mean ± SEM, n=2-6 mice per group. B) A schematic diagram depicts intestinal infection of WT and PINK1 KO mice via gavage at 1- and 2-w.p.i. followed by 3’ scRNAseq of the colon from 2-3 mice per group. C) Multiple cell types identified in the gut of WT and PINK1 KO inf. mice and uninf. control groups at 1- and 2-w.p.i. (in red and blue, respectively) are projected in 2D UMAP plots. Each scRNAseq dataset comprises of ∼40 000 cells and ∼20, 000 genes detected. D) Dotplot indicates the cell identity marker genes curated from the literature used to annotate immune cells in the scRNAseq dataset. E) Merged UMAP plots of immune cell types across all groups at both timepoints. Weeks post-infection, w.p.i.; wild type, WT; knockout, KO; single-cell RNAsequencing, scRNAseq; infected, inf.; uninfected, uninf.; uniform manifold approximation and projection, UMAP.

To gain insights into PINK1-mediated alterations in the intestinal immune landscape, we then performed single cell RNAseq using 10X droplet-based microfluidics technology in FACS-sorted whole colonic lamina propria cells (Fig 1B). Briefly, we prepared gene expression libraries from wild type and PINK1 KO infected mice along with uninfected control groups at 1- and 2-w.p.i. (Fig. 1C). Subsequent to the integration of our datasets employing standard parameters in *Seurat* [27], we annotated all immune cell types using identity markers curated from the literature for each cell cluster (Fig. 1D, E). By comparing wild type and PINK1 KO datasets in each distinct condition (uninfected control, 1-w.p.i. and 2-w.p.i.), we determined transcriptional regulation driven by PINK1 in specific immune cell types in response to intestinal infection. As shown in Fig. 2A, we did not observe obvious genotype-dependent alterations in cell cluster identities in each condition, although as expected, we noticed a larger population of granulocytes at 1-and 2-w.p.i. in both genotypes compared to the uninfected groups. Meanwhile, CD4+ and CD8+ T cell subtypes became more prominent at 2-w.p.i. These changes are consistent with the involvement of innate and adaptive immunity in bacterial clearance at these timepoints [24]. Next, we performed differential gene expression analyses between PINK1 KO and wild type mice. The heatmap displays a number of genes that were either significantly upregulated or downregulated in each immune cell type of PINK1 KO mice (adjusted p-values < 0.05 using non-parametric Wilcoxon rank sum test with Bonferroni correction applied to evaluate significance) with the most substantial changes occurring in the myeloid cell compartment (Fig. 2B). We found extensive transcriptional regulation in monocytes/macrophages and dendritic cells at 1-w.p.i., followed by CD4+ Th1 and Th17 as well as CD8+ cytotoxic T cells (Tc1) at 2-w.p.i.

**Figure 2:**
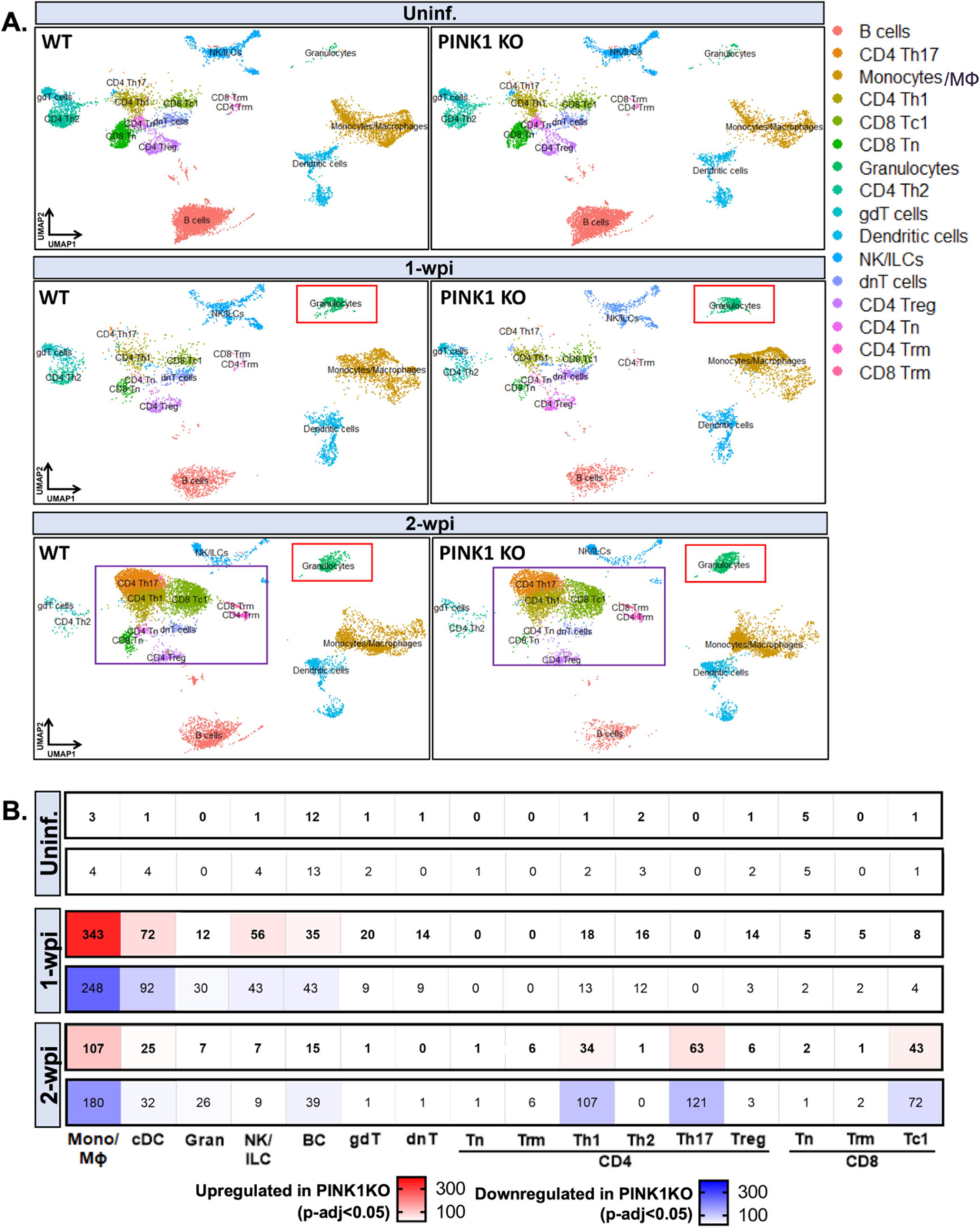
Differential gene expression analyses of the intestinal immune compartment in an infection-induced PD model before the appearance of motor symptoms. A) UMAP plots of immune cell types in the gut segregated by genotype and timepoint. Note cell populations (in boxes) representing a clear expansion of immune cells in both genotypes following infection. B) Each heatmap illustrates the number of significant DEGs in each immune cell types of PINK1 KO compared to WT mice at uninf., 1- and 2-w.p.i. (up-and down-regulated denoted as red and blue, respectively). To test for significance, Wilcoxon Rank Sum test was applied, with adjusted p-value < 0.05 using Bonferroni correction. Helper T cells, Th; cytotoxic 1 T cells, Tc1; naïve T cells, Tn; gamma delta T cells, gdT; natural killer, NK; innate lymphoid cell, ILC; double-negative T cells, dnT; regulatory T cells, Treg; tissue resident memory, Trm; differentially expressed genes, DEGs.

*Higher proportions of effector T cells depicting proinflammatory and cytotoxic characteristics are present at 2-weeks post-infection in the colon of PINK1 KO mice* Effector CD4+ and CD8+ T cell subtypes that appear to be more abundant in peripheral blood, cerebrospinal fluid and brains of PD patients are now widely being investigated due to their proinflammatory and cytotoxic profiles [28-36]. Previous findings by our group also point to the proliferation of autoreactive CD8+ T cells as a prerequisite for driving *late* central nervous system (CNS)-related pathological process via MHCI-mediated antigen presentation by dendritic cells in the spleen [16, 21, 22]. However, it is unknown if PINK1 loss also potentiates the recruitment or expansion of effector T cells and alters their expression profile in the gut at the earliest stages of disease.

To address this, we first examined the features of CD4+ and CD8+ T cells at peak expansion by performing trajectory analyses at 2-w.p.i. using *Slingshot* [37] (Fig. 3A). This bioinformatic tool can identify distinct T cell lineages based on gene expression changes between cell clusters generated by unsupervised hierarchal clustering. Depending on the position in a given trajectory, a pseudotime value is assigned to a cell with increasing number, the further it is from the origin of the lineage which, in this case, was defined as naïve T cells (Tn). In CD4+ T cells, four lineages were obtained starting from Tn towards different effector T cell subtypes: 1) Th1 to helper type 2 (Th2) and IL17-producing Th2 cells; 2) Th1 to resident memory cells (Trm); 3) Th1 to Th17 cells; and 4) Treg (data not shown). Considering that Th1 and Th17 gene expression was most regulated within the CD4+ T cell populations (Fig. 2B), we probed this lineage transition to examine the frequency of these proinflammatory effector T cells (Fig. 3A, upper panel). PINK1 KO mice exhibited higher Tn to effector T cell transitions for both Th1 and Th17 compared to wild type, the genotype-dependent difference in the Th1 transition was most evident with a three-fold higher ratio of Th1-to-Tn (Fig. 3B, upper panel). Pseudotime analyses of CD8+ T cell population revealed a single trajectory from Tn to cytotoxic CD8 Tc1 to Trm cells (Fig. 3A, lower panel). Proportional analyses demonstrated that CD8+ T cell differentiation was greater in PINK1 KO with a four-fold increase in Tc1-to-Tn and Trm-to-Tn ratios compared to wild type (Fig. 3B, lower panel).

**Figure 3:**
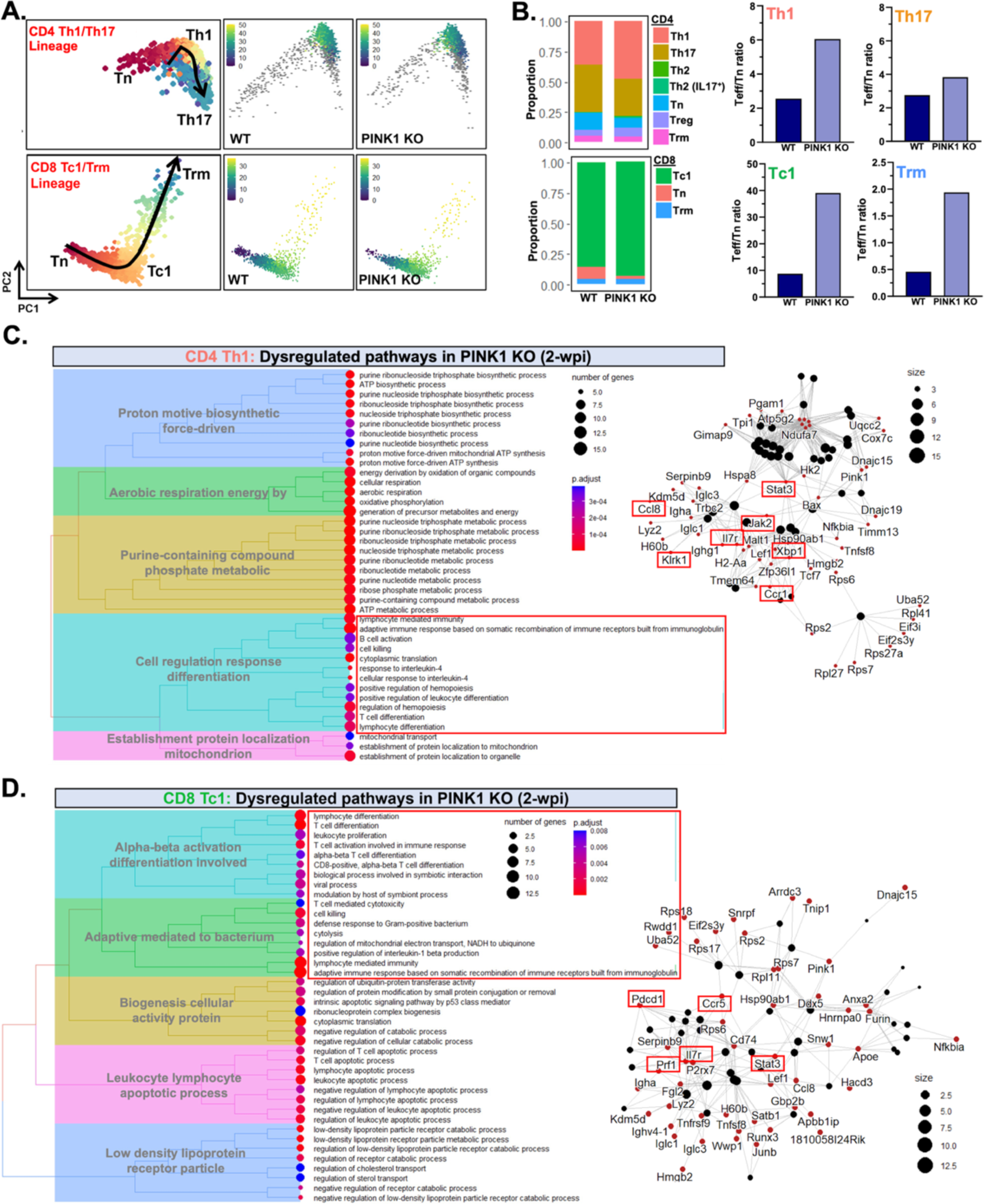
An expansion of proinflammatory and cytotoxic effector T cell subtypes in the gut of PINK1 KO mice following infection. A) Pseudotime analyses of CD4+ and CD8+ T cell subtypes at 2-w.p.i. revealed Th1 to Th17 and Tc1 to Trm lineages from CD4+ and CD8+ Tn, respectively. Both lineages are illustrated in principal component analysis (PCA) plots of intestinal T cell populations from WT and PINK1 KO infected mice. B) The proportions of each CD4+ and CD8+ T cell subtypes were assessed and the ratios of effector-to-naïve T cells (Teff/Tn) in PINK1 KO compared to WT are quantified. C, D) Pathway enrichment analyses were run on significant DEGs in PINK1 KO CD4+ Th1 and CD8+ Tc1 cells against Gene Ontological terms of biological processes database using *ClusterProfiler*. Dysregulated pathways are shown as treeplots while cnetplots display the genes involved, in which the biological processes and associated genes underlying the activation of adaptive immunity are highlighted in rectangles.

Given the expansion of Th1 and Tc1 populations in PINK1KO mice, and the high degree of gene regulation in these populations (Fig. 2B), we conducted pathway enrichment analyses using Gene Ontological (GO) terms to determine the top biological processes that were associated with differentially expressed genes (DEGs) that were significant between PINK1 KO and wild type in each of these T cell subtypes. A large number of GO terms were conserved across T cell types which related to the activation of adaptive immunity (Fig. 3C, 3D). Amongst them were T cell differentiation, cellular response to interleukin-4 (which is imperative in shaping T cell responses), leukocyte cell-cell adhesion, T cell-mediated cytotoxicity and cell killing. Some DEGs that were driving these GO terms (Fig. 3C, 3D) include activation/suppressive markers such as *Stat3*, *Xbp1*, *Cd40lg*, *Il7r*, *Pdcd1*, as well as genes encoding tumor necrosis factor (TNF) ligand superfamilies (*Tnfrsf4*, *Tnfsf8*), C-C chemokine receptors and cytokines (*Ccr1*, *Ccr2*, *Ccr5*, *Ccl8*) and genes involved in cytotoxic T cell function (*Klrk1*, *Klrc1*, *Prf1*). This data suggests that PINK1-mediated immune dysregulation in the gut contributes to the expansion of proinflammatory and cytotoxic effector T cell subtypes early in disease.

### Proinflammatory myeloid cells with enhanced antigen presentation capacity are present at 1-week post-infection in the colon of PINK1 KO mice

In contrast to T cell reactivity, which we postulate to be pivotal for late-stage neurodegeneration [21-23], the role of myeloid cell populations in PD is less clear. About 80% of patients with REM-sleep behaviour disorder (RBD) eventually receive a PD diagnosis [38], making this patient group ideal for studying the prodrome of PD. Studies of the peripheral blood from RBD patients demonstrate abnormal patterns of gene signatures of monocytes which inversely correlate with cognitive/motor performance and functions mediated by midbrain DA neurons in patients [39-41]. This suggests that monocyte features may be an important early biomarker of disease activity. Albeit what monocyte-related molecular pathways might be involved in the initiation of disease is unknown.

Interestingly, our infection-induced PD model indeed substantiates the involvement of monocytes at early stages of disease whereby the highest number of DEGs were found within the myeloid cell lineage, including intestinal monocytes, macrophages (Mϕ) and dendritic cell (DC) populations at 1-w.p.i. (Fig 4A). Specifically, the monocyte population had the highest regulated genes (187 upregulated and 127 downregulated). We delved deeper into the myeloid cell lineage and performed unsupervised clustering on these cells (generating higher resolution principal component analyses) to identify monocytes distinct from macrophages and dendritic cells. We then analyzed the differential trajectories in the myeloid cell lineage using the aforementioned pseudotime methods, providing monocytes as origin given these are the precursor cells for Mϕ and DC (Fig. 4B). In keeping with the literature, we denoted two distinct lineages in both wild type and PINK1 KO mice emerging from monocytes differentiating into either Mϕ or DC. To determine the propensity of intestinal monocytes to differentiate into mature myeloid cells, we quantified the proportions of monocytes from wild type and PINK1 KO mice at 1-w.p.i. with low or high pseudotime values defined as [0, 10] and [10, 20], respectively (given the majority of monocytes fall with the range of 0 to 20) (Fig. 4C). This analysis suggested that PINK1 KO monocytes have a greater tendency to commit to becoming mature myeloid cells as evidenced by a higher ratio of Mϕ or DC-like-to-immature monocytes in PINK1 KO infected mice compared to wild type (Fig. 4C). Most of the top GO terms associated with DEGs in the monocyte population in PINK1 KO infected mice compared to wild type represent functions related to dysregulated mononuclear lymphocyte mediated immunity and myeloid hemopoiesis leukocyte differentiation pathways which are congruent with an altered differentiation state (Fig. 4D). Key DEGs that were driving these GO terms and associated proinflammatory profiles include activation markers such as serum amyloid a3 (*Saa3*), aconitate decarboxylase 1 (*Acod1*), toll-like receptor 2 (*Tlr2*), and hypoxia-inducible factor 1 alpha (*Hif1α*) (Fig 4A, E).

**Figure 4:**
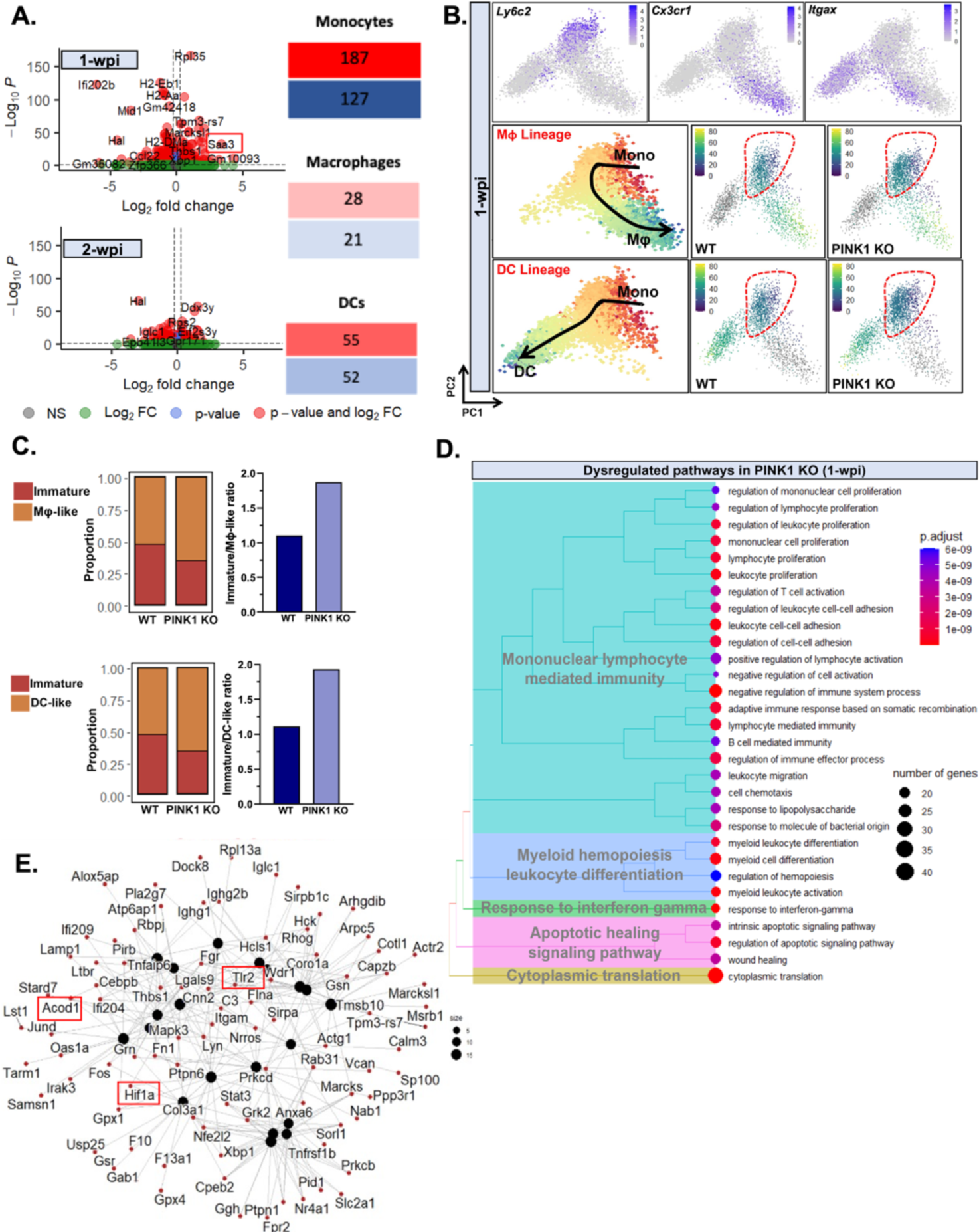
PINK1-driven dysregulation in the myeloid cell lineage at the earliest phase of disease. A) Volcano plots indicating DEGs in PINK1 KO compared to WT of monocytes/macrophages populations at 1- and 2-w.p.i. To test for significance, Wilcoxon Rank Sum test was applied using Bonferroni correction. Significant DEGs (p.adj < 0.5 with a log_2_ fold-change ϕ 0.25) are shown in red. Tables represent the number of DEGs per myeloid cell population whereby upregulated genes are shown in red while downregulated in blue. B) Pseudotime analyses of myeloid cell populations at 1-w.p.i. revealed macrophage (Mϕ) and dendritic cell (DC) lineages originating from monocytes, defined by the expression of *Cx3cr1*, *Itgax* and *Ly6c2*, respectively. Both lineages are illustrated in PCA plots of intestinal myeloid cell populations from WT and PINK1 KO infected mice whereby the monocytes are encircled in red. C) The cells within the monocyte population are classified to either immature or mature (Mϕ-or DC-like) subtypes with pseudotime values of [0, 10] (low) or [10, 20] (high). The proportions of each monocyte subtypes were assessed and the ratios of Mϕ/DC-like-to-immature in PINK1 KO compared to WT are quantified. D, E) Pathway enrichment analyses were run on significant DEGs in monocytes at 1-w.p.i. against Gene Ontological terms of biological processes database using *ClusterProfiler*. Dysregulated pathways are shown as a treeplot while a cnetplot to display the genes involved in driving major GO terms. Boxes represent genes that were subsequently validated using independent approaches.

To corroborate PINK1-driven myeloid cell changes, we isolated CD45^+^CD11b^+^F4/80^-^ Ly6C^high^ monocytes from the colonic lamina propria of PINK1 KO and wild type mice at 1-w.p.i. by fluorescence-activated cell sorting (FACS) along with uninfected control groups (Fig. 5A). We validated significantly altered expression of key genes suggestive of enhanced proinflammatory responses in intestinal monocytes of PINK1 KO infected mice, such as *Lcn2*, *Saa3*, *Acod1*, *Tlr2*, *Hif1α* and a suppression of the anti-inflammatory cytokine, interleukin-10 (*Il10*) (Fig. 5B) which is analogous to critical observations made with single-cell RNAseq (Fig 4A, E). We also noted elevated protein expression of CD11b and CD11c, which are cell surface markers of Mϕ and DC populations, in intestinal Ly6C^high^ monocytes from PINK1 KO infected mice, further attesting to the increased differentiation state of PINK1 KO monocytes following infection (Fig. 5C).

**Figure 5:**
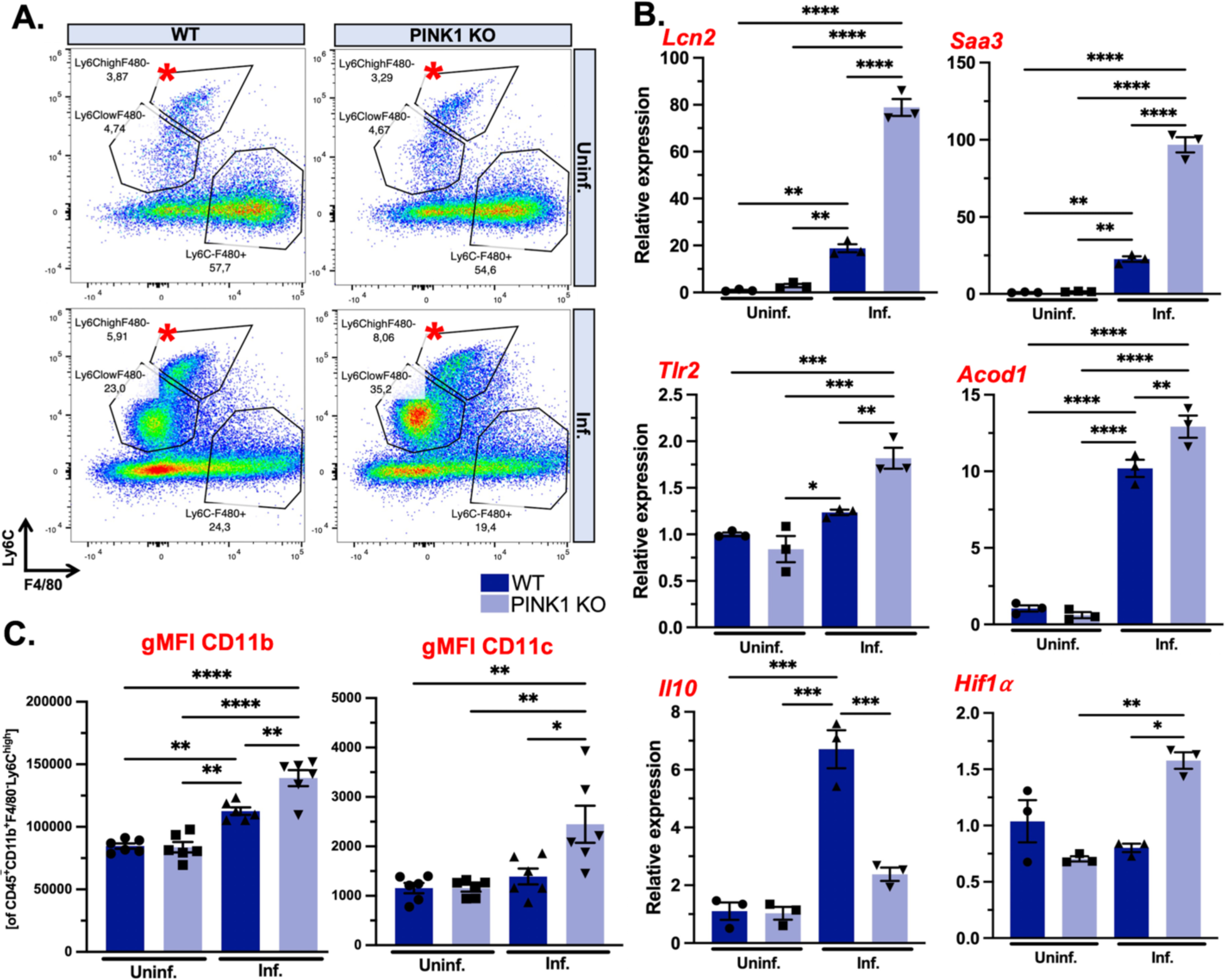
PINK1 KO monocytes acquire enhanced proinflammatory characteristics. A) Representative flow cytometric graph of intestinal myeloid cells (CD45^+^CD11b^+^) from WT and PINK1 KO at 1-w.p.i. Ly6C^high^F4/80^-^ monocytes of intestinal myeloid cells were collected by FACS for downstream gene expression analyses by qPCR. B) Expression of key significant DEGs involved in proinflammatory processes identified in monocytes/macrophages cluster were assessed from PINK1 KO infected mice compared to WT, including *Lcn2*, *Saa3*, *Tlr2*, *Acod1*, *Il10* and *Hif1α*. Two-way ANOVA followed by Tukey’s multiple comparison test was applied. Mean ± SEM, n=3 replicates of 6 pooled mice per group. C) Flow cytometric analyses of Ly6C^high^F4/80^-^ intestinal monocytes for maturation markers were performed by evaluating geometric mean fluorescent intensity of CD11b and CD11c expressed at high levels in macrophages and dendritic cells. Two-way ANOVA followed by Tukey’s multiple comparison test was applied. Mean ± SEM, n=6 mice per group.

### Myeloid and CD8+ T cells are poised for direct cell-cell contact mediated interactions in the colon of infected mice

Previous work has revealed that deregulated antigen presentation can trigger CD8+ T cells that mount a cytotoxic response against mitochondrial self-antigens [16, 21, 23]. Intriguingly, by conducting cell-cell interaction analyses using *CellChat* [42], we bioinformatically uncovered that myeloid and CD8+ T cells have the strongest interactions based on the incoming/outgoing signals received or sent within the immune milieu of the gut (Fig. 6A). Moreover, considering that the probability of the outgoing signaling, which is represented as the outgoing interaction strength, is highest in myeloid cells (monocytes/macrophages and dendritic cells) at both 1- and 2-w.p.i., we assessed their communication to all immune cell types following infection. There were minimal differences in most pathways involved in various cell-cell interactions between wild type and PINK1 KO mice (Fig. 6B). However, we unravelled multiple pathways pertaining to direct cell-cell contact-mediated myeloid cell interactions in both genotypes and revealed that CD8+ Tc1 cells were a major cell type receiving signals. This finding involved signalling pathways related to antigen presentation, such as major histocompatibility complexes (MHC), co-stimulatory molecules (CD86 and CD80), programmed death-ligand 1 (PD-L1) and intercellular adhesion molecule 1 (ICAM1) amongst others highlighted in Fig. 6B. As such, we interrogated the expression of salient molecules in the antigen presentation pathway using flow cytometry of Ly6C^high^ monocytes in the gut following infection of wild type and PINK1 KO mice. These experiments validated elevated levels of MHC, CD86, PDL1 and ICAM1 in PINK1 KO intestinal monocytes, proteins that are also key participants in the interactions between monocytes and CD8+ T cells (Fig. 6C).

**Figure 6:**
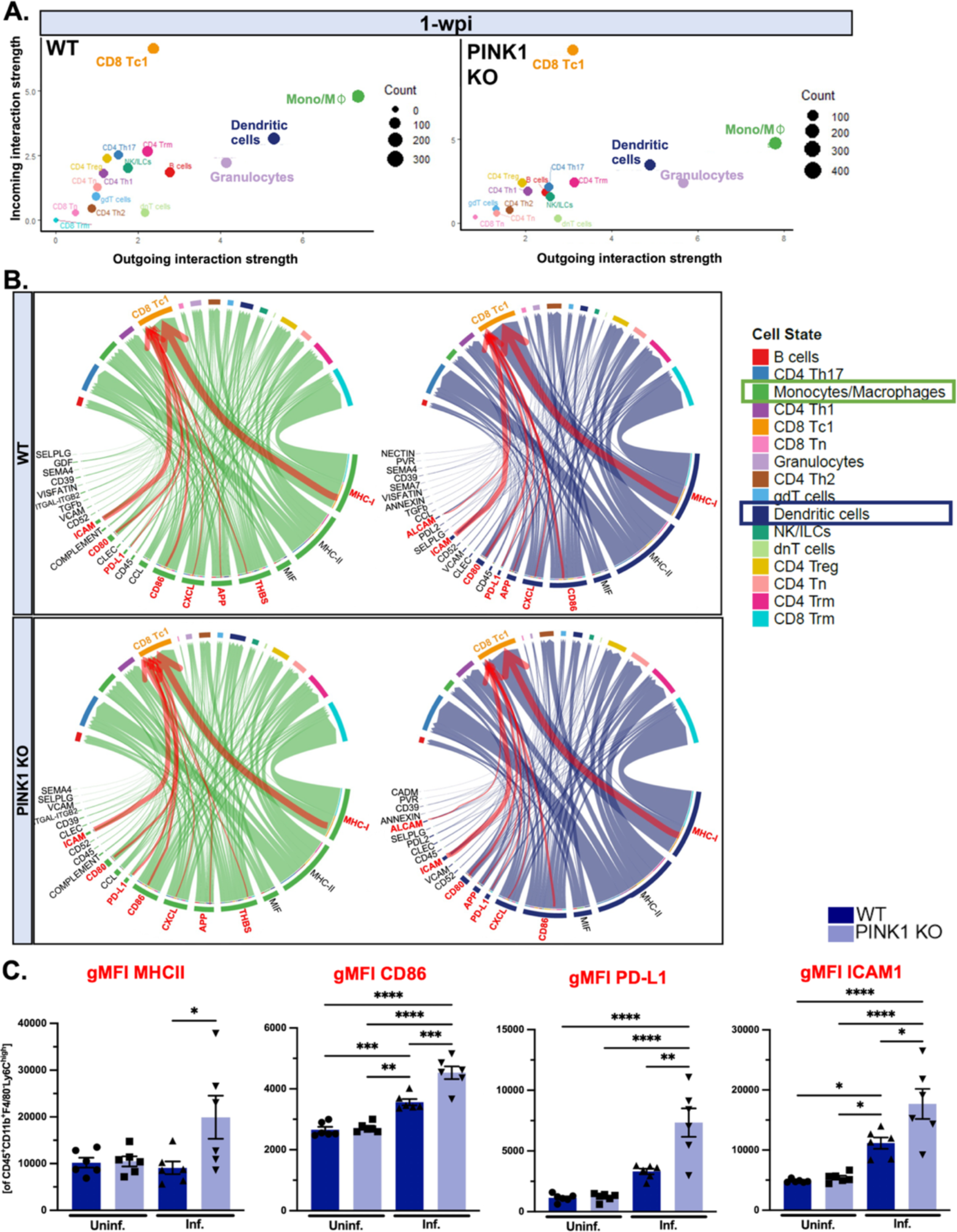
Intercellular communication analyses predict heightened myeloid-cytotoxic CD8+ T cell interactions regulating antigen presentation. B) Unbiased cell-cell interaction analyses of intestinal immune cell populations in WT and PINK1 KO mice at 1- and 2-w.p.i. were conducted. Analysis of incoming and outgoing interaction strengths showed the putative strongest senders and receivers as the myeloid cells (Mono/Mϕ and DC) and effector CD8+ T cells (Tc1 and Trm), respectively. Depicted is a scatter plot of immune cell interactions at 1-w.p.i. B) Chord diagrams illustrate the associations in WT and PINK1 KO monocytes/macrophages (green) and dendritic cells (blue) to neighboring immune cells following infection. Highlighted in red are pathways underpinning myeloid-CD8+ Tc1 cells interaction, which are relevant to antigen presentation. The width of the arrows symbolizes the number of interactions encompassing the specified pathway. C) Flow cytometric analyses of Ly6C^high^F4/80^-^ intestinal monocytes were performed and geometric mean fluorescent intensity of proteins related to antigen presentation and implicated in myeloid-CD8+ T cells interaction were assessed, including MHCII, CD86, PD-L1 and ICAM1. Two-way ANOVA followed by Tukey’s multiple comparison test was applied. Mean ± SEM, n=6 mice per group.

## Discussion

Given the multifaceted nature of PD etiology, it is important to consider the implication of multiple genetic and environmental triggers of this disease, including their impact on immune mechanisms that could drive pathophysiological processes implicated in disease initiation and progression. In the present study, we further characterized a recently established infection model that is geared to study some of the earliest features of PD pathology driven by genetic-environmental interactions. This work unveiled dysregulated signaling pathways in myeloid cells at the initial stages of disease, potentially contributing to antigen presentation and mitochondrial antigen-specific CD8+ T cell-mediated autoimmune mechanisms.

Clinical studies underscore the manifestation of peripheral immune dysfunction in PD wherein inflammatory mediators were reported more abundant in the stools of patients [4-8], indicative of GI inflammation. To decipher the underlying inflammatory-mediated mechanisms at play in the periphery driving pathology, we exploited our previously developed infection-induced PD model before motor symptom onset. Consistent with clinical observations, we found that PINK1 KO infected mice exhibit exacerbated intestinal inflammation as evidenced of higher fecal lipocalin-2 levels compared to wild type. Of note, we inoculated mice with higher *C. rodentium* (4 × 10^-9^ CFU) contrary to our previous publication [21], which a genotype-dependent disparity in inflammatory intestinal marker following infection was not apparent.

In line with a plethora of reports highlighting the role of other genes associated with familial PD in inflammation [43-49], PINK1 has been shown to play pleiotropic effects in innate immunity [17-20, 50]. However, whether PINK1-regulated inflammatory signaling perpetrates early disease processes in PD remains elusive. Here we discovered that the myeloid cell lineage exhibits the most substantial dysregulation at the initial stages of our infection-induced model. We further reveal an amplified proinflammatory response in monocytes of PINK1 KO infected mice, including upregulation of HIF1α. Consistent with this, a prior study demonstrated that deletion of PINK1 in non-immune cells resulted in glucose metabolism reprogramming, which requires increases in reactive oxygen species (ROS) production and the activation of HIF1α signaling [51]. Our data suggest the occurrence of early HIF1α-mediated metabolic alterations in activated PINK1 KO monocytes, which intriguingly is also in line with what has been observed in monocytes from peripheral blood of RBD patients who are considered at the prodromal period of the disease. These patients display elevated levels of mitochondrial ROS associated with a concomitant increase in glycolysis [52]. Other features of RBD monocytes include upregulation of human leukocyte antigen (HLA)-DR expression which correlates with worsened cognitive and motor performance [41], while changes in TLR4 expression inversely correlate with the functionality of midbrain DA neurons, estimated by 18F-DOPA positron emission tomography (PET) imaging [39]. These findings are also recapitulated in our data demonstrating elevated expression of major histocompatibility complexes and toll-like receptors in intestinal monocytes of PINK1 KO mice at the earliest phase of infection.

Cellular roles of PINK1 seems to be context-dependent and operate in response to a distinct type of stressors. Previous studies elucidated conflicting results regarding the implication of PINK1 in mitigating inflammation [17-20, 34]. Sliter *et al.,* reported that PINK1 KO mice under mitochondrial stress display excessive systemic levels of the proinflammatory cytokines interleukin 6 and ϕ-interferon via increased activation of the cyclic GMP–AMP synthase (cGAS)-stimulator of interferon genes (STING) signaling pathway [17]. A previous *in-vitro* study of human skin cells following varicella zoster viral infection also suggested that PINK1 attenuates STING-mediated interferon responses [18]. In contrast, PINK1 was shown to induce retinoic-acid-inducible gene I (RIG-I)-like receptors-mediated antiviral immunity in peritoneal macrophages [19]. Presently, we provide evidence for a negative regulation of myeloid cell responses by PINK1 in an intestinal microbial infection-induced PD model. It is still unclear if this dysregulation in myeloid cells triggered by PINK1 deficiency directly contributes to neuronal dysfunction and loss in the periphery and the CNS or whether the effects are indirect through an activation of cytotoxic T cells. *Sun et al.,* previously reported that deletion of PINK1 in lipopolysaccharide-stimulated microglia elicits the generation of an inflammatory milieu conducive to neuronal death *in vitro* [20]. Understanding the mechanisms for how PINK1 loss potentiates inflammatory responses in myeloid cells and the repercussions on DA neurons will be crucial to develop approaches to target immune-mediated pathological processes implicated in disease onset.

The contribution of PINK1 in inflammation has largely been attributed to its role in regulating mitophagy. During cellular stress, this serine/threonine kinase together with another PD-associated protein, the E3 ubiquitin ligase Parkin, is stabilized at the surface of damaged mitochondria, critical for the initiation of mitophagy. In the absence of PINK1/Parkin, impaired mitophagy occurs and is traditionally viewed as a putative neuronal cell-autonomous mechanism contributing to DA neuron loss [53-55]. Albeit recent work has suggested instead a role of PINK1-driven mitophagy in modulating innate immunity induced by oxidative stress and viral infections [17-19]. Hence, the impact of PINK1 loss of function on neurons could be indirect. Strikingly, our group previously showed that PINK1 represses immune responses independent of mitophagy, through inhibition of the formation of MDVs, which carry proteins to late endosomes, where they are further processed. PINK1 abrogation thus induces MDV transport to the endosomal compartments resulting in heightened MitAP in dendritic cells undergoing cellular stress [16]. With MitAP occurring in the periphery, it is likely that naïve T cells become educated against these self-peptides. Indeed, we have previously revealed that elevated MitAP in PINK1 KO dendritic cells leads to the establishment of peripheral mitochondrial antigen-specific T cells, which can later access the brain and cause degeneration of DA neurons [21-23]. The exact mechanisms linking such processes to neuronal loss are nonetheless unclear. Since DA neurons have the capacity to express MHCI at their cell surface in response to inflammatory signals [21, 56], a direct attack of DA neurons by cytotoxic CD8+ T cells is therefore plausible, as indicated by recent results alluding that adoptive transfer of mitochondria-specific CD8+ T cells in PINK1 KO and wild type mice is sufficient to cause paucity of midbrain DA neurons [23]. Importantly, our current study further showcases that deregulated antigen presentation mediated by PINK1 ablation occurs at the earliest stage of disease as evidenced by upregulation of MHC molecules and co-stimulatory ligands, including CD86 and ICAM1, in intestinal PINK1 KO monocytes. Future work is required to unravel how or if these early aberrant immune responses play a pivotal role in initiating PD-linked autoimmunity.

The present work suggests that following acute *C.rodentium* infection in PINK1 KO mice, myeloid cells in the gut will preferentially interact with CD8+ Tc1 cells via antigen presentation pathways. The activation of intestinal CD8+ Tc1 cells in PINK1 KO mice at early stages after infection is illustrated by upregulation of *Stat3* and *Ccr5*. Prior study demonstrated that CCR5 overexpression induces the activation of the JAK/STAT signaling pathway in human CD4+ T cells [57]. In addition, accumulation of CCR5 at the immunological synapse leads to more robust APC-T cell interactions [58]. Moreover, in this study, CD8+ Tc1 cells in PINK1 KO infected mice displayed elevated levels of cytotoxic mediator perforin (*Prf1*) concomitant with increased expression of *Pdcd1* encoding PD-1, known to be upregulated in antigen-specific T cells upon T-cell receptor engagement. Notably, a related infection model showed that mitochondrial antigen-specific CD8+ T cells express activation-induced marker, PD-1 *in vitro* [22]. Collectively, our findings indicate that not only are effector CD8+ T cells expanding in the gut of PINK1 KO mice at an early stage after intestinal infection, but it is likely that these T cells elicit more potent cytotoxic response. Further characterization of naturally occurring mitochondrial proteins processed and presented through the PINK1-controlled MDV pathway is warranted and will open novel avenues for early detection and interventions in PD.

Ultimately, our study bridges critical knowledge gaps on the influence of genetic and environmental interplay on immune mechanisms that confer risk for PD. A deeper understanding of immunological events underlying the earliest pathological features may permit the development of effective therapies to delay or arrest the progression of disease from the periphery to the CNS.

## Methods

Protocols associated with this work can be found on protocols.io: dx.doi.org/10.17504/protocols.io.kxygxy77ol8j/v1

### Mice

DAT-Ires-Cre (RRID:IMSR_JAX:006660) mice were bred with Ai9 mice (RRID: IMSR_JAX:007909) to induce the expression of tdTomato in DAT-expressing cells (DAT-tdTomato) then were crossed with PINK1-deficient (PINK1 KO) mice in a mixed B6.129 background (RRID:IMSR**_**JAX:017946) [59]. Adult DAT-Cre-tdTomato wild type and PINK1 KO mice (both sexes) were utilized in the present study. Mice were housed under specific pathogen-free conditions, given water and food ad libitum. Studies were approved by the Université de Montréal and McGill University Animal Care and Use Committee (CDEA). CDEA guidelines on the ethical use and care of animals were followed.

### Infection with *C. rodentium*

Mice were infected with chloramphenicol-resistant *C. rodentium* (4 × 10^-9^ CFU) by oral gavage. To monitor burden of *C. rodentium*, faeces were collected from each mouse, weighed, dissociated in PBS, serial-diluted and plated on chloramphenicol-containing MacConkey agar Petri plates. Petri plates were incubated at 37°C overnight to allow colony growth, which were counted the following day. *C. rodentium* was identified by its distinctive morphology on MacConkey agar. Final counts were measured as CFU g^-1^ of faeces. Faecal collection occurred on 1-, 2- and 4-weeks-post-infection (w.p.i.) to confirm 210^7^ CFU/g infection rates (∼80% of infected mice achieved this rate and were included in the study).

### Enzyme-linked immunosorbent assay

Faecal samples were suspended in PBS plus 0.1% Tween20 at a ratio of 100 mg of faeces per mL. To isolate the supernatants, samples were vortexed for 20 mins before centrifugation at 13,000 rpm at 4°C. ELISA was performed using the mouse Lipocalin-2 DuoSet kit (R&D Systems, Cat. No. DY1857) as per manufacturer’s instructions. In brief, collected supernatants were applied to a 96-well plate pre-coated with Lipocalin-2 capture antibody, washed three times with PBS plus 0.05% Tween20, treated with the detection antibody, washed three times, treated with streptavidin–HRP, washed three more times, and developed with a 1:1 ratio of hydrogen peroxide and tetramethylbenzidine. The optical density of the plates was read at 450 nm.

### Colonic lamina propria cell preparation

Following mice euthanasia by CO_2_ asphyxiation and cervical dislocation, colons were dissected then fecal material was flushed with an 18G needle and adipose tissue was removed. Colon was then cut longitudinally and then chopped into sections of 1-2 cm in length. To separate the mucosal epithelium, intestinal tissue sections were incubated in 2 mM EDTA at 37°C for 15 mins followed by vigorous manual shaking three times until cellular suspension appears dense and cloudy. Remaining intestinal tissue sections (consisting of the lamina propria (LP) cells) were then collected and enzymatically dissociated with 2:1 ratio of collagenase IV to DNAse I at 37°C for 30 mins followed by vigorous vortexing. The cellular suspension was then collected through a 70 μm cell strainer. A subsequent similar enzymatic digestion was necessary to completely dissociate intestinal tissue and release the whole lamina propria cells, which was then pooled with the first dissociation. Cells were spun down at 1500 rpm at 4°C for 7 mins at the end of each enzymatic digestion, then the pellet was resuspended in PBS supplemented with 1% fetal bovine serum (FBS). The percentage of viable cells was assessed by flow cytometry; the average was 60%. Cells were subject to downstream experiments, including cell surface staining to acquire cell profile as well as cell sorting for qPCR and single-cell RNAseq.

Once separated, the mucosal epithelium was processed independently for single-cell RNAseq. Briefly, the cloudy cellular suspension containing the crypts was further dissociated into single cells by resuspending in TrypLe using 21G needle followed by 10 mins incubation at 37°C. The reaction was neutralized with basal media (Advanced DMEM F/12 supplemented with 10% FBS). Cellular suspension was then collected through 40 μm cell strainer and spun down at 900 rpm at 4°C for 5 mins. The pellet was resuspended in PBS supplemented with 1% FBS. The percentage of viable cells was assessed by flow cytometry; the average was 40%. Finally, live single cells were collected by FACS.

### Flow cytometry

Following isolation, cells resuspended in PBS were stained using eBioscience fixable viability dye coupled with Zombie Red (Invitrogen, Cat. No. 65-0866-14) for 20 mins on ice, in the dark. Prior to washes with FACS buffer (PBS supplemented with 2% FBS and 1 mM EDTA), cells were spun down at 400 x g at 4°C for 5 mins. Fc receptors were then blocked with anti-mouse CD16/CD32 (Invitrogen, Cat. No. 14-01061-86) along with extracellular staining consisting of a mixture of conjugated antibodies in FACS buffer (1:250) for 30 mins on ice, in the dark. The antibodies used are shown in Table 1. After multiple washes, cells were resuspended in FACS buffer then acquired with LSRFortessa. For sorting live single cells, FACS Aria was used instead. Appropriate controls were applied such as compensation beads and fluorescence minus one (FMOs).

**Table 1:** Extracellular antibodies used for colonic LP staining.

| Antigen | Fluorophore | Clone | Company | Cat. No. | RRID |
| --- | --- | --- | --- | --- | --- |
| CD45 | Brilliant Violet 750 | 30-F11 | BD | 746947 | AB_2871734 |
| CD11b | APC Fire 810 | M1/70 | BioLegend | 101288 | AB_2910274 |
| Ly6C | PerCP-Cy5.5 | HK1.4 | Invitrogen | 45593282 | AB_2723343 |
| F4/80 | APC | BM8 | BioLegend | 123116 | AB_893481 |
| CD11c | PE Fire 810 | QA18A72 | BioLegend | 161106 | AB_2904307 |
| MHC II | APC/Cyanine7 | M5/114.15.2 | BioLegend | 107628 | AB_2069377 |
| CD86 | Alexa Fluor 700 | GL-1 | BioLegend | 105024 | AB_493721 |
| PD-L1 | Brilliant Violet 421 | 10F.9G2 | BioLegend | 124315 | AB_10897097 |
| ICAM1 | FITC | YN1/1.7.4 | BioLegend | 116105 | AB_313696 |

### Quantitative reverse transcription PCR

RNA was extracted using the RNeasy mini kit (Qiagen, Cat. No. 74104) as instructed by the manufacturer. Applied Biosystems^TM^ High-Capacity cDNA Reverse Transcription Kit (FisherScientific, 4368814) was then used for reverse transcription to generate cDNA which was used for real-time quantitative PCR (qPCR) via TaqMan assays (as shown in Table 2) on a QuantStudio^TM^ 5 real-time PCR system. The 2^−ΔΔCt^ method was used to analyse the data using *Actb* as control.

**Table 2:**
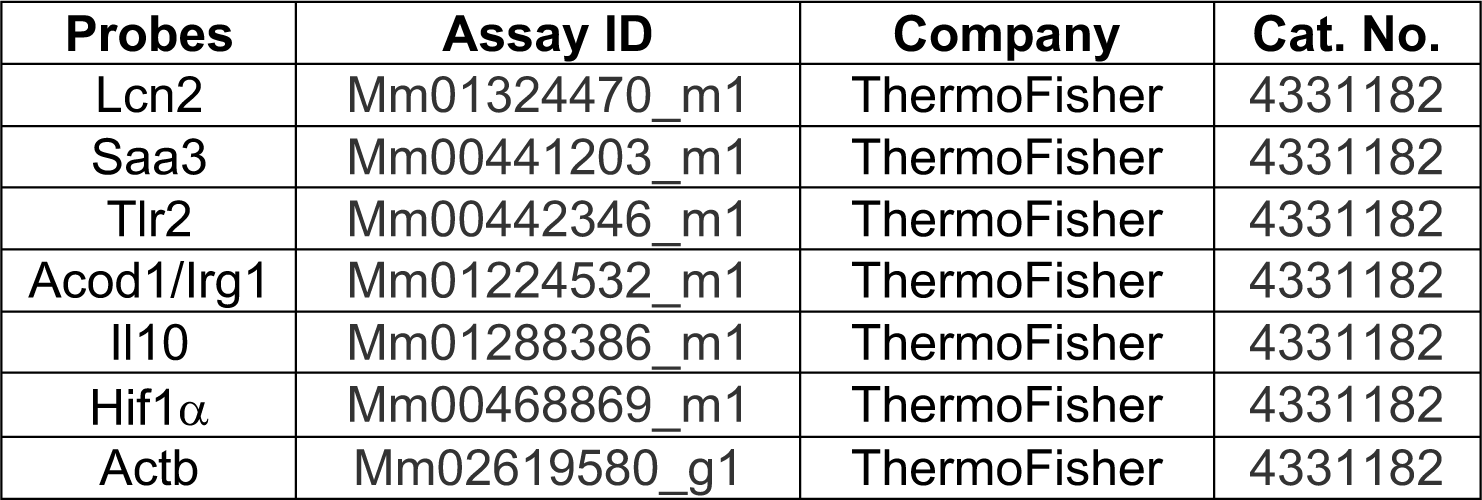
TaqMan gene expression assays used in intestinal monocytes.

### Single-cell library preparation and analysis

Live cells were prepared for single-cell sequencing at final concentration of 1000 cells/μL. All cells were processed according to 10X Genomics Chromium Single Cell 3’ Reagent Guidelines (https://support.10xgenomics.com/single-cell-gene-expression). Briefly, cells were partitioned into nanoliter-scale Gel Bead-In-EMulsions (GEMs) using 10X GemCode Technology. Primers containing (i) an Illumina R1 sequence, (ii) a 16 bp 10x barcode, (iii) a 10 bp Unique Molecular Identifier (UMI) and (iv) a poly-dT primer sequence were incubated with partitioned cells resulting in barcoded, full-length cDNA from poly-adenylated mRNA. Silane magnetic beads were used to remove leftover biochemical reagents/primers, then cDNA was amplified by PCR. Enzymatic fragmentation and size selection was used to optimize cDNA amplicon size prior to library construction. R1 (read 1 primer sequence) were added during GEM incubation, whereas P5, P7, a sample index (i7), and R2 (read 2 primer sequence) were added during library construction via end repair, A-tailing, adaptor ligation and PCR. Quality control and quantification was performed using an Agilent Bioanalyzer High Sensitivity chip. Sequencing was performed using NovaSeq 6000 S4 PE 100bp, which resulted in a read depth of ∼20 000 reads/cell. Reads were processed using the 10X Genomics Cell Ranger Single Cell 2.0.0 pipeline (RRID:SCR_017344, https://support.10xgenomics.com/single-cell-gene-expression/software/pipelines/latest/what-is-cell-ranger) with default and recommended parameters, as previously described [60]. FASTQs generated from sequencing output were aligned to the mouse GRCm38 reference genome using the STAR algorithm 2.7.3a (RRID:SCR_004463, http://code.google.com/p/rna-star/) [61]. Next, gene-barcode matrices were generated for each individual sample by counting unique molecular identifiers (UMIs) and filtering non-cell associated barcodes. This output was then imported into the Seurat v4.9.9.9039 R toolkit v.4.2.1 (for quality control and downstream analysis of our single cell RNAseq experiment [62]. All functions were run with default parameters, unless specified otherwise. Low quality cells (<200 genes/cell and >5 % of mitochondrial genes) were excluded from the overall experiment. Gene expression was log normalized to a scale factor of 10 000.

### Statistical analysis

Statistical analyses were performed using GraphPad Prism 5.0 software. A two-way ANOVA was used to compare the mean of three or more groups of data. *P*-values were adjusted using Tukey’s multiple comparison test. Mean and standard error of the mean (SEM) of biological replicates (*n*) are plotted in all graphs and indicated in the Figure legends. *P-value* < 0.05 was considered statistically significant.

### Illustrations

The graphical abstract and the schematic representation of the experimental designs were created with BioRender.com.

## Acknowledgment

We would like to thank Christina Gavino for the technical support in inoculating mice and all bacterial analyses. We would also like to thank all members of the Desjardins ASAP team for insightful feedback throughout the course of this study. The study is funded by the joint efforts of The Michael J. Fox Foundation for Parkinson’s Research (MJFF) and the Aligning Science Across Parkinson’s (ASAP) initiative. MJFF administers the grant ASAP 000525 on behalf of ASAP and itself. SJR received a CIHR-Vanier Canada Graduate Scholarship. The Trudeau lab also received support from the Canadian Institutes of Health Research (CIHR) (grant PJT-165928).

## Ethics Statement

The animal study was reviewed and approved by the Université de Montréal Animal Care Use Committee (CDEA) and McGill University Animal Care Committee under conditions specified by the Canadian Council on Animal Care and were approved by.

## Author Contributions

SJR led all experiments including cellular, biochemical and bioinformatics work. AK performed flow cytometric staining and helped in the analysis of colonic lamina propria cells. HB performed ELISA experiments. BC and JP helped prepared cells for single cell RNAseq. SP and AM supported biochemical work, including qPCR and flow cytometry. SM bred and maintained animal colony. AA created the website allowing access to and user-friendly analysis of the single cell RNAseq data. EA and MY supported bioinformatics work. LET and HM provided feedback on the project. SJR and JAS prepared figures and drafted the manuscript. SJR, JAS and SG conceived the project, supervised the work, and approved the final manuscript. All authors approved the submitted version.

## Data availability

Raw and processed single cell data available in <u>GSE271210</u> and source code can be found at https://github.com/stratton-lab. Raw tabular data of flow cytometry, qPCR, ELISA and pseudotime values available at https://doi.org/10.5281/zenodo.12041754. Single cell RNAseq data is accessible through a user-interface website available for further analyses (https://singlocell.openscience.mcgill.ca/display?dataset=RNA_Ms_Gut_Immune_PD_2024).

